# Activated RNAi does not rescue piRNA pathway deficiency in testes

**DOI:** 10.1101/2024.07.04.602103

**Authors:** Eliska Taborska, Zuzana Loubalova, Marcos Iuri Roos Kulmann, Radek Malik, Valeria Buccheri, Josef Pasulka, Filip Horvat, Irena Jenickova, Radislav Sedlacek, Petr Svoboda

## Abstract

RNA interference (RNAi) and PIWI-associated RNAs (piRNA) pathways use small RNAs as sequence-specific guides to repress transposable elements. In mice, the loss of *Mili*, an essential piRNA pathway factor, causes male sterility associated with mobilization of LINE L1 retrotransposons while female mutants remain fertile. At the same time, mouse oocytes have exceptionally active RNAi thanks to an oocyte-specific variant of RNase III Dicer, which efficiently makes small RNAs from long dsRNA substrates. In oocytes of mice lacking functional MILI and the oocyte-specific Dicer variant, we previously observed that L1 retrotransposons are redundantly targeted by both, RNAi and piRNA pathways. To test whether enhanced RNAi may reduce the *Mili* mutant phenotype in testes, we used transgenic mice ectopically expressing the oocyte-specific Dicer variant during spermatogenesis. We report here that this genetic modification increases siRNA biogenesis and supports RNAi but is not sufficient to reduce spermatogenic defects caused by the loss of *Mili*.

## Introduction

Protection of the genome against transposable elements (TEs) is vital for maintaining genome integrity across generations. Numerous mechanisms evolved to recognize and suppress TEs, including RNA silencing pathways where small RNAs guide sequence-specific recognition and suppression of TEs. The key RNA silencing pathway suppressing TEs in animal germline is the piRNA pathway (reviewed in (Aravin et al. 2007; Ozata et al. 2019)), which provides adaptive response for silencing active TEs in the genome. The mammalian piRNA pathway is essential for mammalian spermatogenesis (Deng and Lin 2002; Kuramochi-Miyagawa et al. 2004; Carmell et al. 2007; Zheng et al. 2010).

In the female germline, piRNA pathway is essential in hamsters but not in mice (Deng and Lin 2002; Kuramochi-Miyagawa et al. 2004; Carmell et al. 2007; Zheng et al. 2010; Hasuwa et al. 2021; Loubalova et al. 2021; Zhang et al. 2021). Yet, piRNAs were shown to operate during oogenesis where they target retrotransposon RNAs (Lim et al. 2013; Kabayama et al. 2017). Notably, mouse oocytes have also highly active canonical RNAi, a post-transcriptional silencing pathway triggered by dsRNA, which also contributes to TE silencing (Svoboda et al. 2004; Murchison et al. 2007; Tang et al. 2007; Tam et al. 2008; Watanabe et al. 2008). Canonical RNAi in most mammalian cells is inefficient, in part owing to inefficient processing of long dsRNA into siRNAs by RNase III Dicer, which is adapted to process small hairpin precursors of microRNAs (Zhang et al. 2002; Ma et al. 2008; Zapletal et al. 2022; Aderounmu et al. 2023). However, mouse oocytes express a unique Dicer isoform (Dicer^O^), which lacks the HEL1 subdomain of the N-terminal helicase, which has an autoinhibitory effect on long dsRNA processing (Ma et al. 2008). Dicer^O^ efficiently cleaves long dsRNA and supports essential role of canonical RNAi in mouse oocytes (Flemr et al. 2013).

While mouse oocytes have two functional RNA silencing pathways targeting TEs, redundant suppression is TE-specific and concerns especially younger families of L1 and IAP (non-LTR and LTR retrotransposons, respectively). This was shown in mouse mutants lacking one or both pathways: RNAi was eliminated upon deletion of oocyte-specific Dicer^O^ and the piRNA pathway was removed by *Mili* knock-out (Taborska et al. 2019). MILI is a cytoplasmic protein, which generates primary piRNAs and secondary piRNAs by so-called ping-pong mechanism with itself or with MIWI2 (Aravin et al. 2008; Kuramochi-Miyagawa et al. 2008; De Fazio et al. 2011). MILI also cooperates with MIWI2 on silencing of L1 and IAP retrotransposons in mouse fetal testes (Molaro et al. 2014; Manakov et al. 2015) and MILI is important for clearance of retrotransposon transcripts in postnatal testes (De Fazio et al. 2011; Di Giacomo et al. 2013).

We speculated that activation of RNAi during mouse spermatogenesis could reduce severity of the *Mili*^*DAH/DAH*^ phenotype. However, as we report here, expression of Dicer^O^ during spermatogenesis does not reduce severity of the *Mili*^*DAH/DAH*^ phenotype in males.

## Results & Discussion

We sought to generate an animal model with a Cre-inducible Dicer^O^ variant to selectively activate RNAi *in vivo*. To that end, we have produced a transgene Tg(EGFP-lox66-pCAG-lox71i-**Dicer**^**O-HA**^-T2A-mCherry), for simplicity referred to as *Dicer*^*Tg(O-HA)*^ hereafter (Fig. 1A). The transgene was designed for inducible expression of Dicer^O-HA^ protein, a C-terminally HA-tagged Dicer^O^ variant, and mCherry reporter. Dicer^O-HA^ was separated from the mCherry protein by a T2A self-cleaving peptide (Ryan et al. 1991). Expression was driven by the CAG promoter, a strong synthetic promoter used for high expression in mice (Miyazaki et al. 1989; Okabe et al. 1997). The transgenic vector also carried homologous arms for ROSA26 locus insertion and two heterologous loxP sites for CRE recombinase (Araki et al. 2010) flanking the CAG promoter oriented away from *Dicer*^*O-HA*^ coding sequence. In this configuration, the promoter can be inverted by the CRE recombinase to activate Dicer^O-HA^ expression (Fig. 1B). The cloned transgene and insertion into ROSA26 in ESC showed that the promoter can flip and activate expression in the opposite direction upon transient transfection of a Cre-expressing plasmid (Fig. 1C-F). While the efficiency of Dicer^O-HA^ expression activation was low, presumably due to poor transfection efficiency of ESCs, these data implied that transgene’s design works. Thus, we have used two ESC clones to establish a transgenic mouse line, which was successfully achieved with one of the clones (Fig. 1G). Testing of Cre-mediated transgene activation suggested that overexpression of Dicer^O-HA^ is detrimental for the survival of mice, as we did not obtain any progeny with activated transgene expression when using *Sox*-CRE and *ActB*-CRE drivers (Fig. 1H). At the same time, we discovered that the uninduced transgene in the ROSA26 locus had leaky expression, which varied across organs, with the highest expression observed in testes (Fig. 1I).

**Figure 1.**
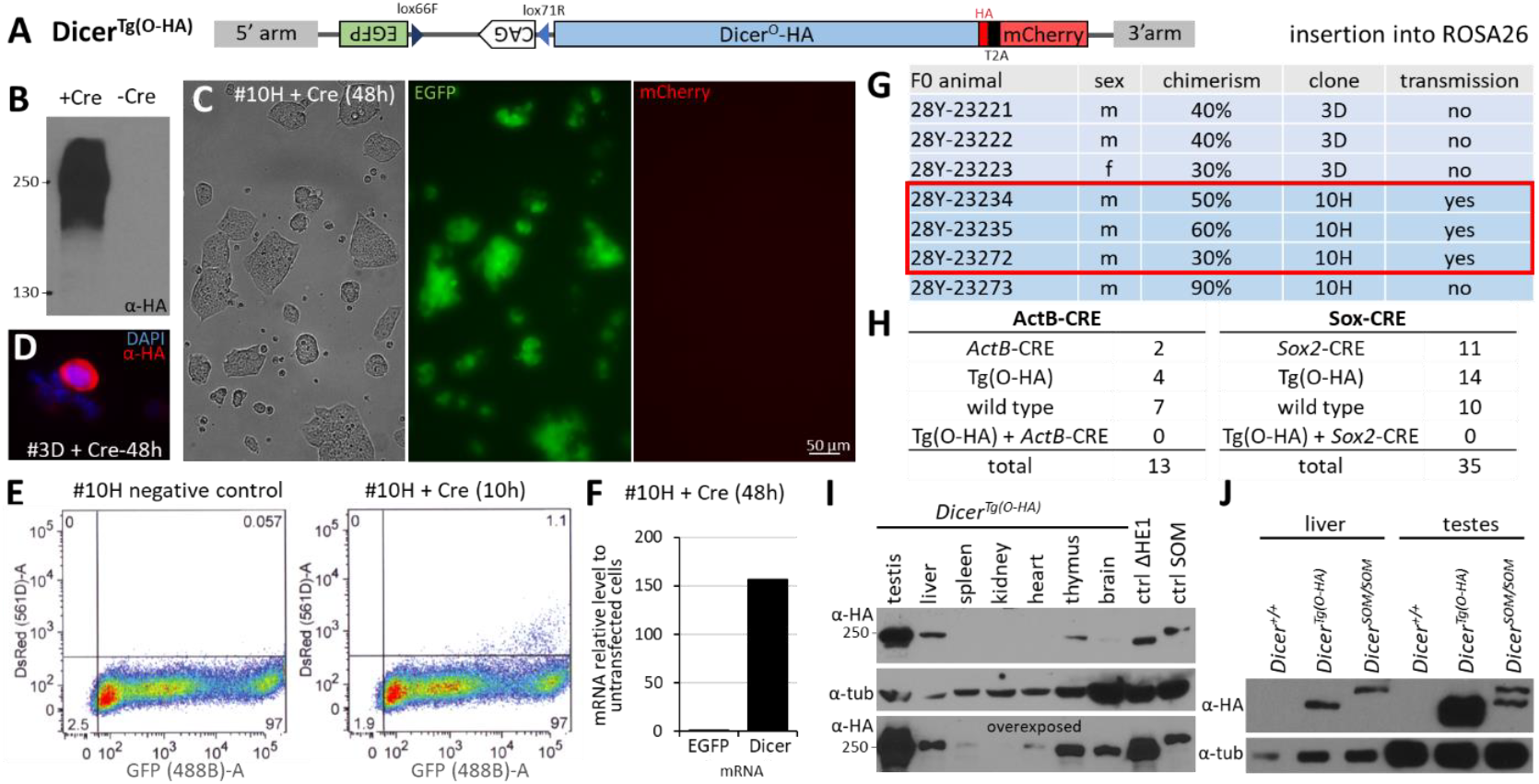
Inducible *Dicer*^*Tg(O-HA)*^ transgene. (A) A scheme of the inducible Dicer transgene. (B) Dicer^O-HA^ expression in transient co-transfection of a transgene plasmid with and without a Cre-expressing plasmid. Dicer^O-HA^ was detected using an α-HA antibody. (C) EFGP and mCherry expression in the transgenic ESC clone #10H 48 hours after transfection of a Cre-expressing plasmid. (D) A rare HA-positive cell in the transgenic ESC clone #3D 48 hours after transfection of a Cre-expressing plasmid. (E) FACS analysis of Cre-induced expression of mCherry in the clone #10H. (F) qPCR analysis of EFGP and *Dicer*^*Tg(O-HA)*^ RNA levels in the transgenic ESC clone #10H 48 hours after transfection of a Cre-expressing plasmid. RNA levels are shown relative to untransfected clone #10H cells. (G) Overview of F0 chimeric animals among 23 live born pups. The three framed founders transmitted the transgene into the next generation. (H) Ubiquitous transgene activation appears lethal – no mice were born when Dicer^O-HA^ expression was induced with two different Cre-drivers. (I) Despite the CAG promoter is oriented into the opposite direction than Dicer expression, the transgene has leaky expression even without induction, which varies among organs, testis being the organ with the highest leakage. (J) Comparison of endogenous HA-tagged Dicer expression (*Dicer*^*SOM/SOM*^) and leaky HA-tagged transgene expression (Dicer^Tg(O-HA)^) in liver and testes suggests that the transgene produces an equivalent of endogenous Dicer amount in liver while in in testes it is strongly overexpressed. The nature of the lower band in the *Dicer*^*SOM/SOM*^ lane is unclear, possibly it is *Dicer*^*Tg(O-HA)*^ loading contamination since *Dicer*^*SOM*^ allele expression yielded only a single band in testes (Zapletal et al. 2022).

To estimate the amount of leaky transgene expression, we compared Dicer^O-HA^ expression with expression of full-length Dicer tagged in the endogenous locus with the HA tag (SOM allele, described in (Taborska et al. 2019)). In the liver, the leaky expression of *Dicer*^*Tg(O-HA)*^ appeared equivalent to that of the endogenous full-length Dicer while in the testis Dicer^O-HA^ was strongly overexpressed (Fig. 1J). The leaky expression of Dicer^O-HA^ protein was also confirmed by high level of mCherry fluorescence in testis of *Dicer*^*Tg(O-HA)*^ animals (Fig. 2A). HA epitope staining in histological sections revealed cytoplasmic signal in spermatogenic cells, including pre-meiotoic, meiotic and post-meiotic cells (Fig. 2B). The highest signal appeared in elongated spermatids but other spermatogenic cells in seminiferous tubules exhibited well-detectable signal (Fig. 2B).

**Figure 2.**
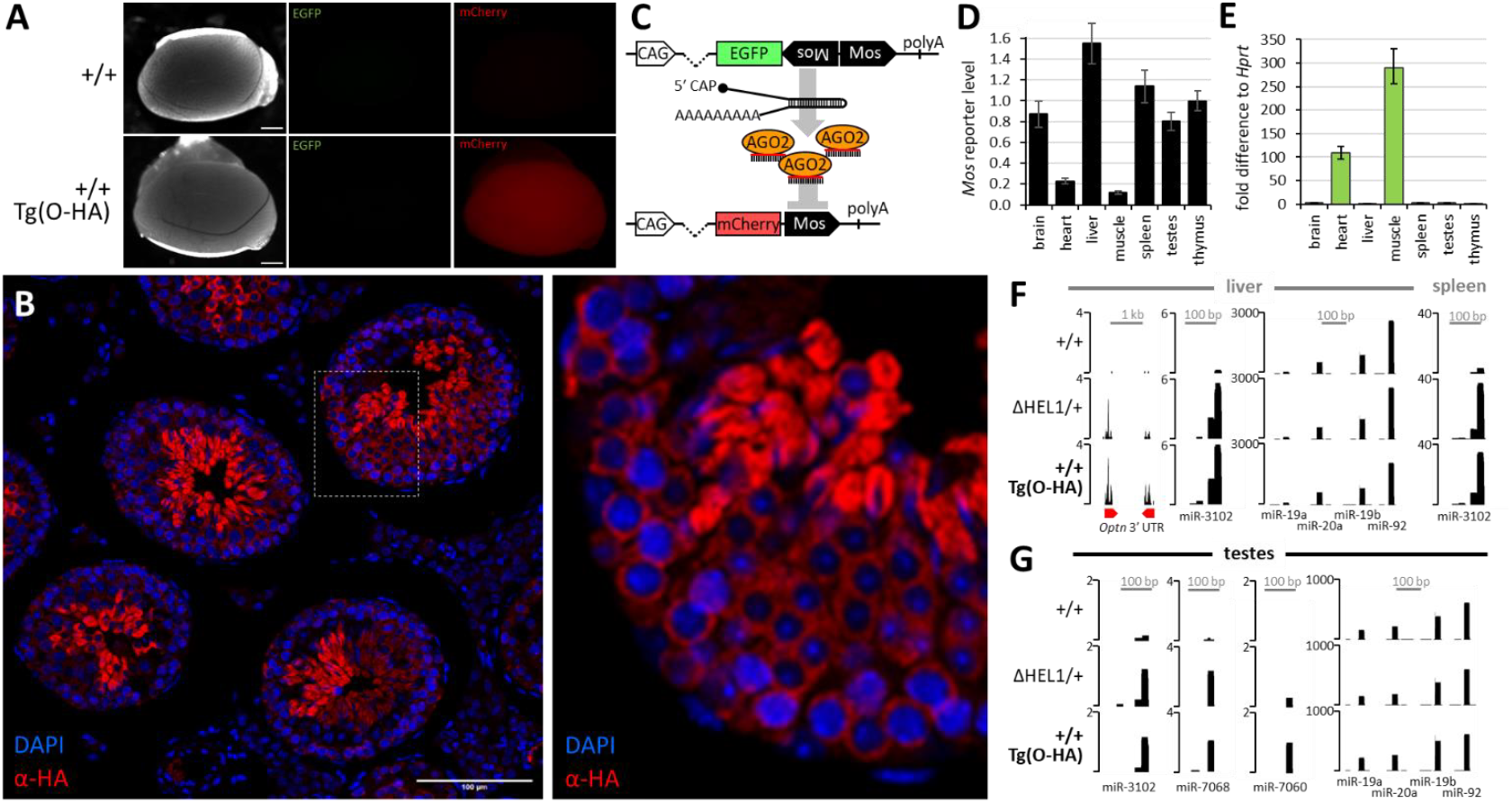
Effects of leaky expression of the *Dicer*^*Tg(O-HA)*^ transgene. (A) Leaky expression in testes yields detectable levels of mCherry reporter from the transgene. (B) mCherry from the Dicer^Tg(O-HA)^transgene appears to be most expressed in post-meiotic cells but it is well detectable in pre-meiotic and meiotic cells as well. Red = α-HA staining, blue = DAPI, size bar = 100 μm. (C) A scheme of the transgenic RNAi system using the transgene Tg(CAG-EGFP-MosIR) as a source of dsRNA and the transgene Tg(CAG-mCherry-Mos) as a sequence-specific target. (D) qPCR analysis of RNAi-targeted reporter in Dicer^Tg(O-HA)^ mice carrying CAG-EGFP-MosIR transgene relative to control mice lacking CAG-EGFP-MosIR. (E) qPCR analysis of CAG-EGFP-MosIR transgene expression in *Dicer*^*SOM/wt*^ *Pkr*^*–/–*^ organs from (Buccheri 2024) for comparison with (D). Expression is presented as fold difference relative to *Hprt*. Data come from three (muscle and spleen) or five (other organs) biological replicates analyzed in technical triplicates (Buccheri 2024). (F) The Dicer^Tg(O-HA)^ transgene increases *Optn* endo-siRNA levels in liver and mirtron levels in liver and spleen. (G) The Dicer^Tg(O-HA)^ transgene increases mirtron levels in testes. Vertical scale depicts counts per million (CPM) of 19-32 nt reads.

The cause of the apparent switch in transcriptional direction of the transgene (EGFP was not detectable in testes, Fig. 2A) is unclear. The integrated transgene in ESCs was expressing EGFP but not mCherry as designed (Fig. 1C), yet after genome reprogramming events in the germline cycle *in vivo*, transcriptional direction of the transgene became reversed. Furthermore, the leaky transcription does not follow the established expression pattern of the CAG promoter (Okabe et al. 1997), which we have observed in several independent transgenic lines (Nejepinska et al. 2012; Buccheri 2024), indicating activation of a cryptic promoter.

We tested whether the leaky expression of *Dicer*^*Tg(O-HA)*^ increases siRNA biogenesis and RNAi. For RNAi, we have used a system (Fig. 2C) where a MosIR transgene is expressing long dsRNA hairpin and a targeted reporter carries a complementary sequence in its 3’UTR (Nejepinska et al. 2012; Buccheri 2024). This system was recently used to analyze RNAi in mice where one endogenous Dicer allele was converted to Dicer^O^-like variant by removing exons encoding the HEL1 subdomain (*Dicer*^*ΔHEL1*^ allele) (Buccheri 2024). Similarly, we produced mice carrying *Dicer*^*Tg(O-HA)*^, *MosIR*, and *Mos* reporter transgenes, and compared *Mos* reporter levels in mice carrying the *Dicer*^*Tg(O-HA)*^ transgene in the presence or absence of the *MosIR* transgene. We have observed strong RNAi effect in heart and muscle in the presence of the *MosIR* transgene (Fig. 2D,E), which is consistent with effects observed in *Dicer*^*ΔHEL1/+*^ mice (Buccheri 2024). At the same time, absence of RNAi in other organs, which are expressing higher levels of the truncated Dicer variants (Fig. 1I and 2D), is consistent with MosIR transgene expression being a limiting factor for RNAi (Buccheri 2024). In any case, these data show that the *Dicer*^*Tg(O-HA)*^ transgene produces highly active Dicer variant able to support RNAi in the heart and muscles, even when low-expressed.

To obtain evidence that siRNA production is enhanced across tissues, we sequenced small RNAs from liver, spleen and testes and analyzed levels of small RNAs previously shown to be upregulated by the Dicer^ΔHEL1^ variant. *Optn* and *Anks3* loci are harboring inverted repeats with potential to produce long dsRNA upon transcription (Tam et al. 2008; Watanabe et al. 2008; Flemr et al. 2013). *Optn*-derived 21-23 nt small RNA levels were low but apparently upregulated in liver in *Dicer*^*Tg(O-HA)*^ animals comparably to *Dicer*^*ΔHEL1/+*^ mice (Fig. 2F). *Optn* and *Anks3* loci in spleen and testis were not expressed at sufficient levels to produce endo-siRNAs. We thus analyzed mirtrons, a class of non-canonical miRNAs, whose precursors were shown to be suboptimal substrates for the full-length Dicer but were efficiently cleaved by the Dicer^ΔHEL1^ variant (Zapletal et al. 2022; Buccheri 2024). The miR-3102 precursor is a prominent non-canonical miRNA example, having a longer stem carrying a miRNA tandem, which is cleaved twice by Dicer. We have observed increased levels of miR-3102 in *Dicer*^*Tg(O-HA)*^ liver while canonical miRNAs did not exhibit such a change (Fig. 2F). The same effect was observed in spleen (Fig. 2F) and testes (Fig. 2G). Additional miR-7060 and miR-7068, two mirtrons previously found to be upregulated in the presence of the Dicer^ΔHEL1^ variant (Zapletal et al. 2022; Buccheri 2024), had also increased levels in testis (Fig. 2G). These data suggest that the *Dicer*^*Tg(O-HA)*^ transgene supports efficient siRNA production also in testis.

Finally, we tested whether the expression of the *Dicer*^*Tg(O-HA)*^ transgene in spermatogenic cells could reduce the *Mili* mutant phenotype. We previously observed that RNAi and piRNA pathways in mouse oocytes redundantly target the same L1 and IAP subfamilies, which are also expressed in spermatogenic cells (Taborska et al. 2019) and upregulated in *Mili* mutants (De Fazio et al. 2011; Di Giacomo et al. 2013; Molaro et al. 2014; Manakov et al. 2015). In addition, RNA-seq suggested these L1 subfamilies produce increased amounts of 21-23 nt siRNAs but not 24-32 nt piRNAs in *Dicer*^*Tg(O-HA)*^ testes (Fig. 3A). We thus hypothesized that activation of RNAi via Dicer^O^ expression might provide suppression of L1 and IAP retrotransposons, which might compensate some effects of the loss of the piRNA pathway. Accordingly, we used *Mili*^*DAH*^ mutant animals (De Fazio et al. 2011), which were used to examine piRNA and RNAi redundancy in oocytes (Taborska et al. 2019), crossed them with *Dicer*^*Tg(O-HA)*^ animals, and analyzed testicular histology of *Mili*^*DAH/DAH*^ in the presence and absence of Dicer^O^. However, these results showed that the Dicer^O^ expression does not improve spermatogenesis defects present in *Mili*^*DAH/DAH*^ mutants (Fig. 3B).

**Figure 3.**
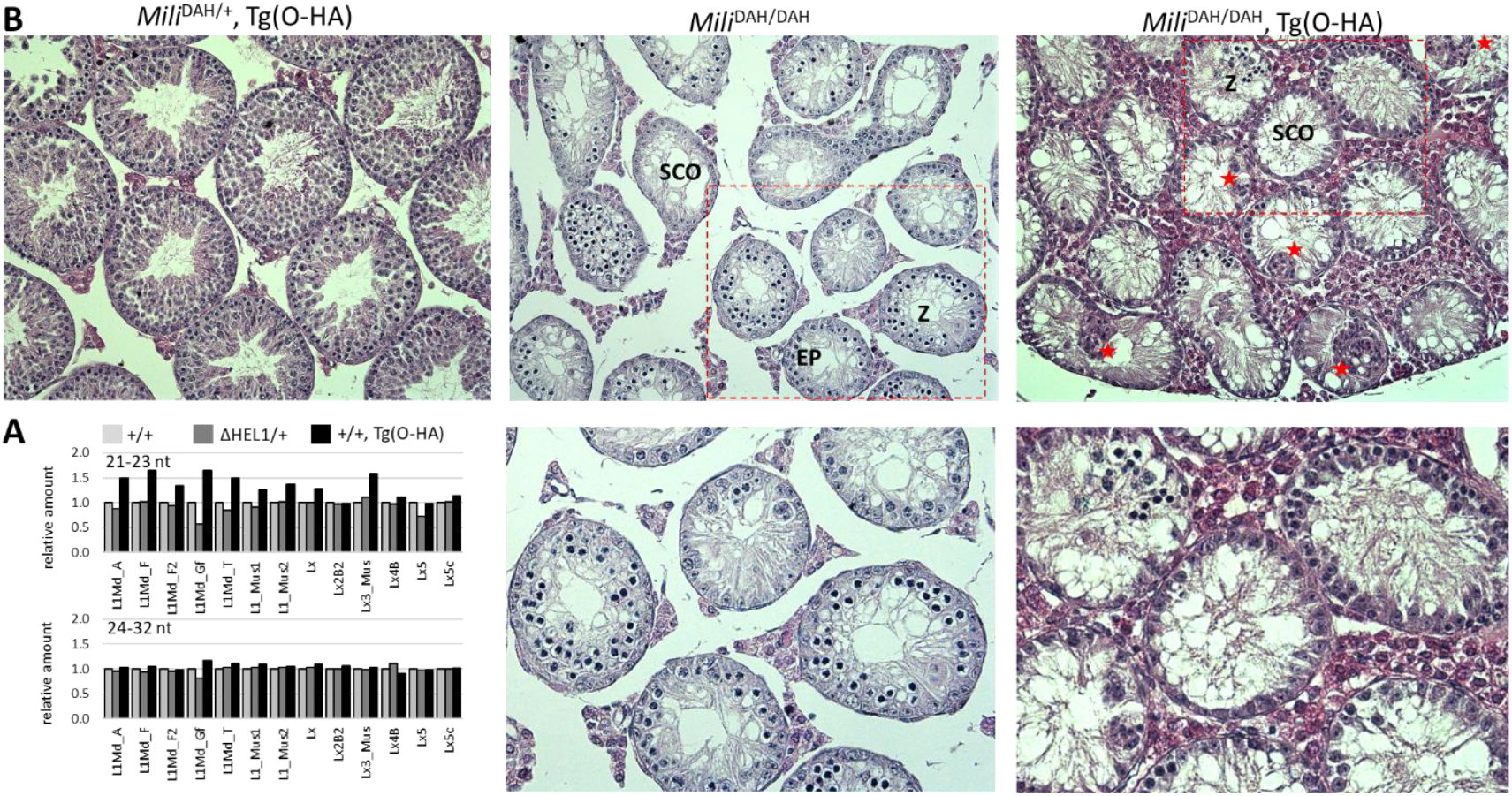
Expression of truncated Dicer^O^ variant yields more LINE L1 siRNAs in testes but it does not reduce the *Mili* mutant phenotype. (A) Quantification of putative siRNA (21-23 nt) and piRNA (24-23) nt reads in RNA-seq data from normal C57Bl/6 (+/+), *Dicer*^*ΔHEL1/+*^ (ΔHEL1/+) and transgenic (+/+, Tg(O-HA)) animals. Abundance of reads perfectly mapping to L1 element subfamilies with full-length intact elements analyzed in oocytes (Taborska et al. 2019) was calculated per 1 million 19-23 reads for each sample and is shown relative to +/+, which was set to one. Data were obtained from age-matched mice, one animal per genotype. (B) Histological sections of 10 weeks-old mice. Dicer^Tg(O-HA)^transgene heterozygote mice on the *Mili*^*DAH/+*^ background show normal seminiferous tubule histology. As previously described (De Fazio et al. 2011), *Mili*^*DAH/DAH*^ mice show the same phenotype as *Mili*^−/−^ with early pachytene as the most advanced stage of germ cell development. Zygotene is the most common stage found in tubules, and a fraction of tubules represents Sertoli-cell only appearance. The *Dicer*^*Tg(O-HA)*^ transgene heterozygote on the *Mili*^*DAH/DAH*^ background shows tubules containing zygotene spermatocytes, however the presence of Sertoli-cell only tubules is more pronounced than in *Mili*^*DAH/DAH*^ mouse. In addition, the tubules show frequent occurrence of cell clusters (giant cell formation with eosinophilic cytoplasm and an indistinct cell border) projecting into the tubular lumen (red asterisks). These observations suggest an additional defect beyond that previously described in mice lacking active MILI, attributed to the presence of the *Dicer*^*Tg(O-HA)*^ allele. Z – zygotene, EP – early pachytene, SCO – Sertoli cells only.

## Conclusions

Taken together, in the mouse female germline the piRNA pathway is nonessential and shows redundancy with RNAi in targeting L1 and IAP retrotransposons. On the other hand, in testes, when RNAi is activated by the same mechanism as in oocytes, there is no observable functional redundancy with the piRNA pathway, which would reduce severity of the *Mili*^*DAH/DAH*^ phenotype.

## Materials and Methods

### Animals

Animal experiments were carried out in accordance with the Czech law and were approved by the Institutional Animal Use and Care Committee (approval no. 34-2014). Tg(EGFP-lox66-pCX-lox71i-Dicer^O-HA^-IRES-mCherry) mice were generated from genetically modified ESCs in the Czech Centre for Phenogenomics. Briefly, the transgene was inserted by homologous recombination into the ROSA26 locus in ESCs; the insertion was facilitated with a CRISPR-Cas9 vector (Flemr and Buhler 2015). Selected ESC clones were injected into morulae and seven chimeric mice were obtained from 233 transferred embryos. The selected line was then bred for at least five generations with normal C57Bl/6NCrl mice.

#### Genotyping

For genotyping, tail biopsies were lysed in DEP-25 DNA Extraction buffer (Top-Bio) according to manufacturer’s instructions. 1 µl aliquots were mixed with primers and Combi PPP Master Mix (Top-Bio) for genotyping PCR. Genotyping primers are listed in Table S1.

#### Organ harvesting

Organs collected from sacrificed animals were either directly used for analysis or snap frozen and stored at -80° C for later use.

#### Tissue histology

Testes from 10-week-old mice were fixed in Bouin’s fixative (Sigma-Aldrich, HT10132-1L) for 2 hours, washed several times with ice-cold PBS, and stored in 70% ethanol at 4°C until processing. Tissues were embedded in paraffin, sectioned at 6 μm thickness on a microtome, mounted from warm distilled water (42°C) onto silane-coated microscope slides, and dried overnight at 42°C. The tissues were then stained with hematoxylin solution (Gill No. 2) for 1.5 minutes, washed with running water, and stained with Eosin for 30 seconds. Slides were mounted with xylene-based mounting medium and coverslips.

### Immunofluorescence analysis of histological sections

Testes were fixed in Hartman’s fixative (Sigma-Aldrich, H0290) overnight at 4°C. Tissues were dehydrated in ethanol, embedded in paraffin, sectioned to a thickness of 2.5–6 μm and used for immunofluorescence staining. Sections were deparaffinized and then boiled for 18 min in 10 mM pH 6 sodium citrate solution for antigen retrieval. After 45 min blocking with 5% normal donkey serum and 5% bovine serum albumin (BSA) in PBS, sections were incubated for 1 h at room temperature or overnight at 4 °C with the primary antibodies used at 1:1000 dilution. Nuclei were stained with 1 μg ml−1 DAPI for 10 min, slides were mounted in ProLong Diamond Antifade Mountant (Thermo Fisher Scientific) and images were acquired using the Leica SP8 confocal microscope.

### Cell culture

Mouse ESCs were cultured in 2i-LIF media: KO-DMEM (Gibco) supplemented with 15% fetal calf serum (Sigma), 1x L-Glutamine (Thermo Fisher Scientific), 1x non-essential amino acids (Thermo Fisher Scientific), 50 µM β-Mercaptoethanol (Gibco), 1000 U/mL LIF (Isokine), 1 µM PD0325901, 3 µM CHIR99021(Selleck Chemicals), penicillin (100 U/mL), and streptomycin (100 µg/mL).

### Transfection

For transfection, ES cells were plated on gelatin-couated 24-well plate, grown to 80% density and transfected using Lipofectamine 3000 (Thermo Fisher Scientific) according to the manufacturer’s protocol. The total amount of transfected plasmid DNA was 1 μg/well. Cells were collected for analysis 48 hours post-transfection.

### Western blot

Mouse organs and ES cells were homogenized mechanically in RIPA lysis buffer supplemented with 2x protease inhibitor cocktail set (Millipore) and loaded with SDS dye. Protein concentration was measured by Bradford assay (Bio-Rad) and 80-100 μg of total protein was used per lane. Proteins were separated on 5.5% polyacrylamide (PAA) gel and transferred on PVDF membrane (Millipore) using semi-dry blotting for 60 min, 35 V. The membrane was blocked in 5% skim milk in TBS-T, Dicer was detected using anti-HA 3F10 monoclonal primary antibody (High Affinity rat IgG1, Roche #11867431001; dilution 1:500), anti-HA rabbit primary antibody (Cell Signaling, #3724, dilution 1:1,000) and incubated overnight at 4°C. Secondary anti-Rat antibody (Goat anti-Rat IgG, HRP conjugate, ThermoFisher #31470, dilution 1:50,000), HRP-conjugated anti-Mouse IgG binding protein (Santa-Cruz #sc-525409, dilution 1:50,000) or anti-Rabbit-HRP antibody (Santa-Cruz #sc-2357, dilution 1:50,000) was incubated 1 h at room temperature. For TUBA4A detection, samples were separated on 10% PAA gel and incubated overnight at 4 °C with anti-Tubulin (Sigma, #T6074, dilution 1:10,000). HRP-conjugated anti-mouse IgG binding protein (Santa-Cruz, #sc-525409, dilution 1:50,000) was used for detection. Signal was developed on films (X-ray film Blue, Cole-Parmer #21700-03) using SuperSignal West Femto Chemiluminescent Substrate (Thermo Scientific).

### RT-qPCR analysis

Cultured cells or mouse organs were harvested, washed in PBS, and homogenized in Qiazol lysis reagent (Qiagen); total RNA was isolated by phenol–chloroform extraction according to the manufacturer’s protocol. 1 µg of total RNA was reverse transcribed using LunaScript RT SuperMix Kit (New England Biolabs) according to the manufacturer’s instructions and 1 µl cDNA was used as a template for a 10 µl qPCR reaction. qPCR was performed on LightCycler 480 (Roche) and the Maxima SYBR Green qPCR master mix (Thermo Fisher Scientific) was used for the qPCR reaction. qPCR was performed in technical triplicates for each biological sample. Average Ct values of the technical replicates were normalized to the housekeeping genes *Hprt, B2m* and *Alas1* using the ΔΔCt method (Pfaffl et al. 2002). qPCR primers are listed in Table S1.

### Small RNA sequencing (RNA-seq) and bioinformatic analyses

Small RNA sequencing from testes and its analysis was done as described previously (Buccheri 2024). Libraries were sequenced by 75-nucleotide single-end reading using the Illumina NextSeq500/550 platform. Data were deposited in the Gene Expression Omnibus database under accession ID GSE267971.

## Acknowledgements

Main funding was provided by the Czech Science Foundation EXPRO grant 20-03950X. Development of genetically modified Dicer mouse models was funded previously from the European Research Council under the European Union’s Horizon 2020 research and innovation programme (grant agreement No 647403, D-FENS). Financial support of M.K. and V.B. was in part provided by the Charles University in a form of a PhD student fellowship; this work will be in part used to fulfil requirements for a PhD degree and hence can be considered “school work”. Institutional support was provided by the Ministry of Education, Youth, and Sports (MEYS) project NPU1 LO1419. The authors used services of the Czech Centre for Phenogenomics at the Institute of Molecular Genetics supported by the Czech Academy of Sciences RVO 68378050 and by the project LM2018126 and LM2023036 Czech Centre for Phenogenomics provided by MEYS. We also acknowledge services of the Light Microscopy Core Facility, IMG, Prague, Czech Republic, supported by MEYS – LM2023050 and RVO – 68378050-KAV-NPUI. Computational resources were provided by the e-INFRA CZ project (ID:90254), supported by the Ministry of Education, Youth and Sports of the Czech Republic and by the ELIXIR-CZ project (ID:90255), part of the international ELIXIR infrastructure.

## Disclosure and Competing Interests Statement

Authors declare no competing interests.

**Table S1.**
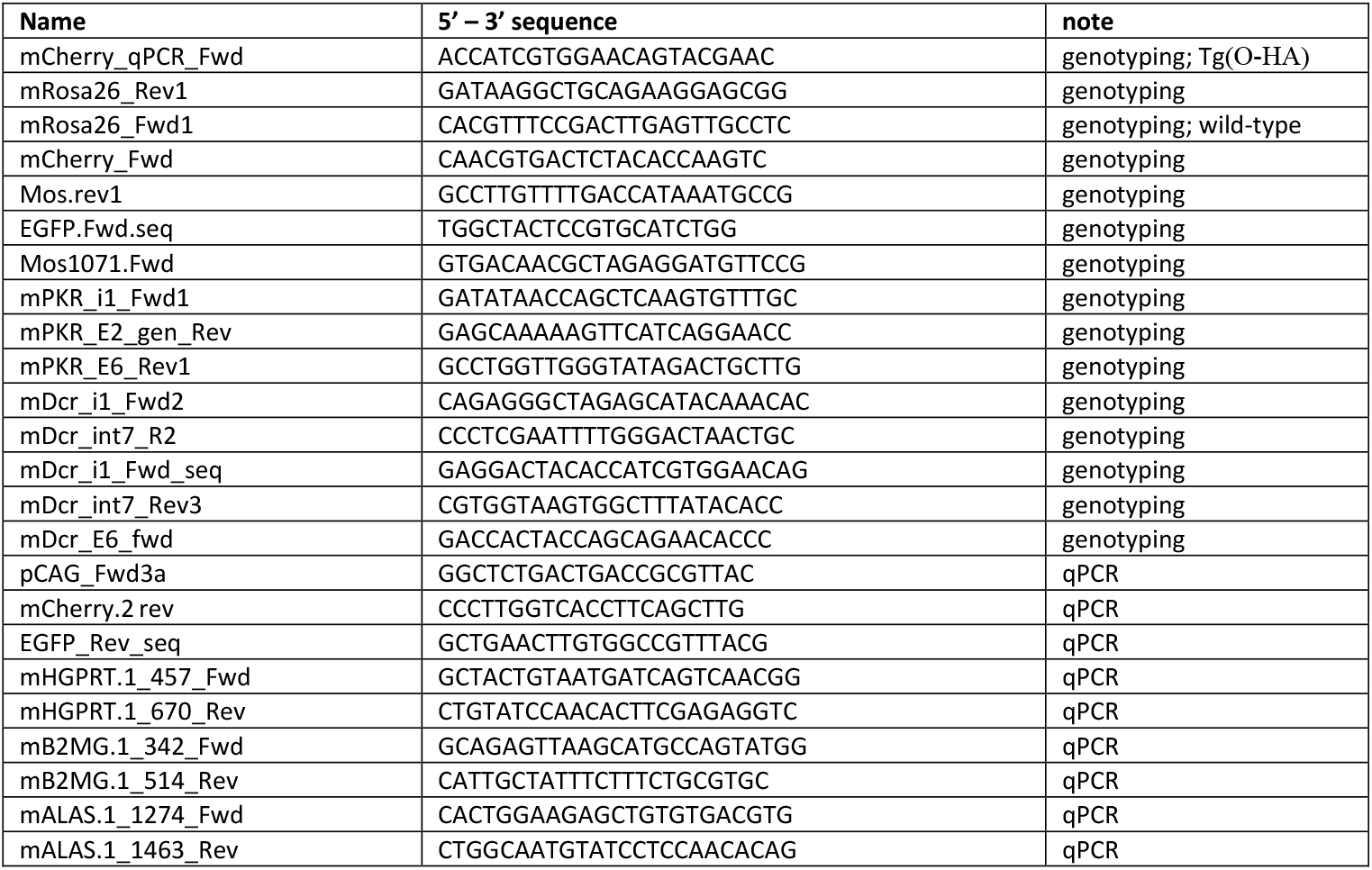
Primers.

## References

Aderounmu AM, Aruscavage PJ, Kolaczkowski B, Bass BL. 2023. Ancestral protein reconstruction reveals evolutionary events governing variation in Dicer helicase function. Elife 12.

Araki K, Okada Y, Araki M, Yamamura K. 2010. Comparative analysis of right element mutant lox sites on recombination efficiency in embryonic stem cells. BMC Biotechnol 10: 29.

Aravin AA, Hannon GJ, Brennecke J. 2007. The Piwi-piRNA pathway provides an adaptive defense in the transposon arms race. Science 318: 761–764.

Aravin AA, Sachidanandam R, Bourc’his D, Schaefer C, Pezic D, Toth KF, Bestor T, Hannon GJ. 2008. A piRNA pathway primed by individual transposons is linked to de novo DNA methylation in mice. Molecular Cell 31: 785–799.

Buccheri Vea. 2024. Functional canonical RNAi in mice expressing a truncated Dicer isoform and long dsRNA. EMBO Reports in press.

Carmell MA, Girard A, van de Kant HJG, Bourc’his D, Bestor TH, de Rooij DG, Hannon GJ. 2007. MIWI2 is essential for spermatogenesis and repression of transposons in the mouse male germline. Developmental Cell 12: 503–514.

De Fazio S, Bartonicek N, Di Giacomo M, Abreu-Goodger C, Sankar A, Funaya C, Antony C, Moreira PN, Enright AJ, O’Carroll D. 2011. The endonuclease activity of Mili fuels piRNA amplification that silences LINE1 elements. Nature 480: 259–263.

Deng W, Lin HF. 2002. miwi, a murine homolog of piwi, encodes a cytoplasmic protein essential for spermatogenesis. Developmental Cell 2: 819–830.

Di Giacomo M, Comazzetto S, Saini H, De Fazio S, Carrieri C, Morgan M, Vasiliauskaite L, Benes V, Enright AJ, O’Carroll D. 2013. Multiple Epigenetic Mechanisms and the piRNA Pathway Enforce LINE1 Silencing during Adult Spermatogenesis. Molecular Cell 50: 601–608.

Flemr M, Buhler M. 2015. Single-Step Generation of Conditional Knockout Mouse Embryonic Stem Cells. Cell Rep 12: 709–716.

Flemr M, Malik R, Franke V, Nejepinska J, Sedlacek R, Vlahovicek K, Svoboda P. 2013. A retrotransposon-driven Dicer isoform directs endogenous small interfering RNA production in mouse oocytes. Cell 155: 807–816.

Hasuwa H, Iwasaki YW, Au Yeung WK, Ishino K, Masuda H, Sasaki H, Siomi H. 2021. Production of functional oocytes requires maternally expressed PIWI genes and piRNAs in golden hamsters. Nat Cell Biol 23: 1002–1012.

Kabayama Y, Toh H, Katanaya A, Sakurai T, Chuma S, Kuramochi-Miyagawa S, Saga Y, Nakano T, Sasaki H. 2017. Roles of MIWI, MILI and PLD6 in small RNA regulation in mouse growing oocytes. Nucleic Acids Research 45: 5387–5398.

Kuramochi-Miyagawa S, Kimura T, Ijiri TW, Isobe T, Asada N, Fujita Y, Ikawa M, Iwai N, Okabe M, Deng W et al. 2004. Mili, a mammalian member of piwi family gene, is essential for spermatogenesis. Development 131: 839–849.

Kuramochi-Miyagawa S, Watanabe T, Gotoh K, Totoki Y, Toyoda A, Ikawa M, Asada N, Kojima K, Yamaguchi Y, Ijiri TW et al. 2008. DNA methylation of retrotransposon genes is regulated by Piwi family members MILI and MIWI2 in murine fetal testes. Genes Dev 22: 908–917.

Lim AK, Lorthongpanich C, Chew TG, Tan CW, Shue YT, Balu S, Gounko N, Kuramochi-Miyagawa S, Matzuk MM, Chuma S et al. 2013. The nuage mediates retrotransposon silencing in mouse primordial ovarian follicles. Development 140: 3819–3825.

Loubalova Z, Fulka H, Horvat F, Pasulka J, Malik R, Hirose M, Ogura A, Svoboda P. 2021. Formation of spermatogonia and fertile oocytes in golden hamsters requires piRNAs. Nat Cell Biol 23: 992–1001.

Ma E, MacRae IJ, Kirsch JF, Doudna JA. 2008. Autoinhibition of human dicer by its internal helicase domain. J Mol Biol 380: 237–243.

Manakov S, Pezic D, Marinov G, Pastor W, Sachidanandam R, Aravin A. 2015. MIWI2 and MILI Have Differential Effects on piRNA Biogenesis and DNA Methylation. Cell Reports 12: 1234–1243.

Miyazaki J, Takaki S, Araki K, Tashiro F, Tominaga A, Takatsu K, Yamamura K. 1989. Expression vector system based on the chicken beta-actin promoter directs efficient production of interleukin-5. Gene 79: 269–277.

Molaro A, Falciatori I, Hodges E, Aravin AA, Marran K, Rafii S, McCombie WR, Smith AD, Hannon GJ. 2014. Two waves of de novo methylation during mouse germ cell development. Genes & Development 28: 1544–1549.

Murchison EP, Stein P, Xuan Z, Pan H, Zhang MQ, Schultz RM, Hannon GJ. 2007. Critical roles for Dicer in the female germline. Genes Dev 21: 682–693.

Nejepinska J, Malik R, Filkowski J, Flemr M, Filipowicz W, Svoboda P. 2012. dsRNA expression in the mouse elicits RNAi in oocytes and low adenosine deamination in somatic cells. Nucleic Acids Res 40: 399–413.

Okabe M, Ikawa M, Kominami K, Nakanishi T, Nishimune Y. 1997. ‘Green mice’ as a source of ubiquitous green cells. FEBS letters 407: 313–319.

Ozata DM, Gainetdinov I, Zoch A, O’Carroll D, Zamore PD. 2019. PIWI-interacting RNAs: small RNAs with big functions. Nat Rev Genet 20: 89–108.

Pfaffl MW, Horgan GW, Dempfle L. 2002. Relative expression software tool (REST) for group-wise comparison and statistical analysis of relative expression results in real-time PCR. Nucleic Acids Res 30: e36.

Ryan MD, King AM, Thomas GP. 1991. Cleavage of foot-and-mouth disease virus polyprotein is mediated by residues located within a 19 amino acid sequence. J Gen Virol 72 (Pt 11): 2727–2732.

Svoboda P, Stein P, Anger M, Bernstein E, Hannon GJ, Schultz RM. 2004. RNAi and expression of retrotransposons MuERV-L and IAP in preimplantation mouse embryos. Dev Biol 269: 276–285.

Taborska E, Pasulka J, Malik R, Horvat F, Jenickova I, Jelic Matosevic Z, Svoboda P. 2019. Restricted and non-essential redundancy of RNAi and piRNA pathways in mouse oocytes. PLoS Genet 15: e1008261.

Tam OH, Aravin AA, Stein P, Girard A, Murchison EP, Cheloufi S, Hodges E, Anger M, Sachidanandam R, Schultz RM et al. 2008. Pseudogene-derived small interfering RNAs regulate gene expression in mouse oocytes. Nature 453: 534–538.

Tang F, Kaneda M, O’Carroll D, Hajkova P, Barton SC, Sun YA, Lee C, Tarakhovsky A, Lao K, Surani MA. 2007. Maternal microRNAs are essential for mouse zygotic development. Genes Dev 21: 644–648.

Watanabe T, Totoki Y, Toyoda A, Kaneda M, Kuramochi-Miyagawa S, Obata Y, Chiba H, Kohara Y, Kono T, Nakano T et al. 2008. Endogenous siRNAs from naturally formed dsRNAs regulate transcripts in mouse oocytes. Nature 453: 539–543.

Zapletal D, Taborska E, Pasulka J, Malik R, Kubicek K, Zanova M, Much C, Sebesta M, Buccheri V, Horvat F et al. 2022. Structural and functional basis of mammalian microRNA biogenesis by Dicer. Mol Cell 82: 4064–4079 e4013.

Zhang H, Kolb FA, Brondani V, Billy E, Filipowicz W. 2002. Human Dicer preferentially cleaves dsRNAs at their termini without a requirement for ATP. EMBO J 21: 5875–5885.

Zhang H, Zhang F, Chen Q, Li M, Lv X, Xiao Y, Zhang Z, Hou L, Lai Y, Zhang Y et al. 2021. The piRNA pathway is essential for generating functional oocytes in golden hamsters. Nat Cell Biol 23: 1013–1022.

Zheng K, Xiol J, Reuter M, Eckardt S, Leu NA, McLaughlin KJ, Stark A, Sachidanandam R, Pillai RS, Wang PJ. 2010. Mouse MOV10L1 associates with Piwi proteins and is an essential component of the Piwi-interacting RNA (piRNA) pathway. Proceedings of the National Academy of Sciences of the United States of America 107: 11841–11846.

